# Ultra-conserved sequences in the genomes of highly diverse *Anopheles* mosquitoes, with implications for malaria vector control

**DOI:** 10.1101/2021.01.13.426530

**Authors:** Samantha M. O’Loughlin, Annie J. Forster, Silke Fuchs, Tania Dottorini, Tony Nolan, Andrea Crisanti, Austin Burt

## Abstract

DNA sequences that are exactly conserved over long evolutionary time scales have been observed in a variety of taxa. Such sequences are likely under strong functional constraint and they have been useful in the field of comparative genomics for identifying genome regions with regulatory function. A potential new application for these ultra-conserved elements has emerged in the development of gene drives to control mosquito populations. Many gene drives work by recognising and inserting at a specific target sequence in the genome, often imposing a reproductive load as a consequence. They can therefore select for target sequence variants that provide resistance to the drive. Focusing on highly conserved, highly constrained sequences lowers the probability that variant, gene drive-resistant alleles can be tolerated.

Here we search for conserved sequences of 18bp and over in an alignment of 21 *Anopheles* genomes, spanning an evolutionary timescale of 100 million years, and characterise the resulting sequences according to their location and function. Over 8000 ultra-conserved elements were found across the alignment, with a maximum length of 164 bp. Length-corrected gene ontology analysis revealed that genes containing *Anopheles* ultra-conserved elements were over-represented in categories with structural or nucleotide binding functions. Known insect transcription factor binding sites were found in 48% of intergenic *Anopheles* ultra-conserved elements. When we looked at the genome sequences of 1142 wild-caught mosquitoes we found that 15% of the *Anopheles* ultra-conserved elements contained no polymorphisms. Our list of *Anopheles* ultra-conserved elements should provide a valuable starting point for the selection and testing of new targets for gene-drive modification in the mosquitoes that transmit malaria.

## INTRODUCTION

DNA sequences that are highly conserved over long evolutionary timescales have been identified in many organisms. Some of these sequences show complete conservation at the nucleotide level and are often known as ultra-conserved elements (UCEs). Originally, UCEs were defined as sequences of at least 200bp that were identical between human, mouse and rat genomes (Bejerano *et al*. 2004). Subsequently the search for UCEs has been extended to other vertebrates, insects and plants (e.g. Makunin *et al*. 2013; Siepel *et al*. 2005; Baxter *et al*. 2012), and to sequences of length 50bp or more.

There are several reasons why UCEs are of interest. First, in the field of comparative genomics, UCEs are thought to represent functionally important regions. While there is still some mystery around why sequences might be conserved at the nucleotide level over long evolutionary timescales, it has been shown that UCEs 1) often are involved in regulation of transcription of genes, especially essential genes involved in development (e.g. Visel *et al*. 2008); 2) may have a role in chromosomal structure (e.g. Chiang *et al*. 2008); and 3) are sometimes non-coding RNA genes (e.g. Kern *et al*. 2015). Even UCEs in protein coding regions may have multi-functional roles (Warnefors *et al*. 2016). Second, UCEs can act as probes to facilitate genomic sequencing of non-model organisms using sequence-capture methods (Faircloth *et al*. 2012). Third, alterations in UCEs have been shown to have an association with human cancers (e.g. Calin *et al*. 2007; Lin *et al*. 2012).

A new potential role for UCEs has recently emerged in the fight against malaria using gene-drive mosquitoes (Kyrou *et al*. 2018). *Anopheles* mosquitoes are the vectors of malaria parasites, and mosquito control has been responsible for much of the recent success in reduction of malaria cases (78% of the 663 million malaria cases averted globally since 2000 (Bhatt *et al*. 2015)). Progress in reducing malaria cases has stalled (WHO 2018), probably in part due to resistance of the mosquitoes against commonly used pesticides. One novel method under consideration is the development of mosquitoes containing gene drives that either reduce the population size (Windbichler *et al*. 2011; Hammond *et al*. 2016) or make them unable to transmit the malaria parasite (Gantz *et al*. 2015). Both methods currently rely on nuclease-based synthetic gene drive systems that introduce a desired trait at a precise genomic location, spreading it in a target population at such a rate that outweighs fitness costs associated with the trait (Burt 2003). The technologies include RNA-guided endonucleases (such as CRISPR/Cas9) and homing endonucleases (Jinek *et al*. 2012; Windbichler *et al*. 2011). These enzymes recognise and cleave a particular target size of about 18 bp. When the sequence coding for these enzymes is engineered into its own target site in the genome and is expressed in the germline, it creates a double-strand break in the homologous chromosome. The break will usually be repaired by homology-directed repair using the drive-containing chromosome as a template which results in conversion of the repaired to also contain the drive element in greater than the usual 50% inheritance rate among the gametes. An efficient gene drive can be inherited by almost 100% of progeny (Hammond *et al*. 2015). Theoretical and laboratory studies have shown that changes to the recognition site can result in alleles that cannot be recognised or cleaved. If these alleles confer increased fitness compared to the wild type allele in the presence of the gene drive they can be expected to spread and retard the spread of the gene drive (Deredec *et al*. 2008; Unckless *et al*. 2017; Hammond *et al*. 2017). For population suppression gene drives that are designed to impair essential genes the selection pressure for resistance alleles to arise is high. These alleles can arise from standing variation at the target site in a wild population, or may come about from the action of the endonuclease. This is because non-homologous end joining can sometimes repair the double-strand break, and random insertions and deletions can be introduced to the target site.

Two of the most important vector species in sub-Saharan Africa are the close relatives *Anopheles gambiae* and *An. coluzzii*, both of which are highly genetically diverse. A study of 765 mosquitoes in phase 1 of Ag1000G project, which looked to sample genetic diversity in the wild through the resequencing of wild caught individuals across Africa (*Anopheles gambiae* 1000 Genomes Consortium 2017), found a polymorphism on average every 2.2 bases of the accessible genome. Nucleotide diversity (π) ranged from ~0.008 to ~0.015 per population sampled, and even non-degenerate sites (which are expected to be strongly constrained) had an average π of ~0.0025.

Proof of principle for retarding the evolution of resistance to nuclease-based gene-drive by targeting an evolutionarily conserved sequence has recently been demonstrated. A strain of mosquitoes with a CRISPR/Cas9 gene-drive targeting the *doublesex* gene fully suppressed laboratory caged populations of *An. gambiae* (Kyrou *et al*. 2018) without selecting for resistance. The CRISPR/Cas9 target sequence in this strain is an intron/exon junction that is highly conserved across the *An. gambiae* species complex, and only one rare single nucleotide polymorphism was found in the sequence in *An. gambiae* and *An. coluzzii* in the Ag1000G data. Consistent with the target site being a region of high functional constraint, monitoring of potential resistant mutations during the cage experiment revealed that although some indels had been introduced by the endonuclease, none of them showed signs of positive selection.

This strong constraint at the nucleotide level may exist at other loci in *An. gambiae*. The Ag1000G project looked for conserved putative CRISPR/Cas9 target sites (18 invariant bases followed by the −NGG motif necessary for Cas9 cleavage) in the 765 mosquitoes of Phase 1 of the project, and found 5474 genes containing such sequences. However, they note that more variation is likely to be found with further sampling.

Here we take an approach that is likely to be more stringent in identifying functionally constrained sequences by searching for regions that are ultra-conserved across the whole *Anopheles* genus, which has a most recent common ancestor ~100 million years ago (Neafsey *et al*. 2015). Although sequence constraint across such a long time scale is not necessary for a good target (as indicated by the *doublesex* locus, which is ultra-conserved within the *An. gambiae* species complex, but shows less conservation outside the complex), we are hypothesising that such highly conserved sequences will contain few polymorphisms in the wild *Anopheles gambiae* population, and any polymorphisms that do arise (either spontaneously or due to the action of the endonuclease) are likely to have strong fitness costs. We also do not confine our analysis to sequences compatible with any single nuclease architecture (e.g the 5’-NGG-3’ PAM sequence required by the SpCas9 nuclease) since the range and flexibility of nuclease architectures is constantly expanding, meaning that these requirements may be relaxed (Anders *et al*. 2016; Chaterjee *et al*. 2018; Hu *et al*. 2018). We extracted UCEs from an alignment of the genomes of 21 *Anopheles* species and strains that was constructed by the *Anopheles* 16 genomes consortium (Neafsey *et al*. 2015). We characterise the UCEs according to their locations in the genome, and use data from *Drosophila* orthologues to group genic UCEs according to potential phenotype. We then use the Ag1000G data (765 *An. gambiae* and *An. coluzzii*) to see whether these conserved elements contain any variation in natural populations of potential target mosquito species.

## METHODS

### Data

Two sources of genomic data were used in this study: a multi-species alignment from the *Anopheles* 16 genomes project (Neafsey *et al*. 2015) and variation data from phase 1 of the MalariaGEN *An. gambiae* 1000 genomes project (Anopheles gambiae 1000 Genomes Consortium 2017). The *Anopheles* 16 genomes project multi-species alignment contains reference genomes from 21 *Anopheles* species and strains: *An. gambiae PEST, An. gambiae s.s., An. coluzzii, An. merus, An. arabiensis, An. quadriannulatus, An. melas, An. christyi, An. epiroticus, An. minimus, An. culicifaces, An. funestus, An. stephensi S1, An. stephensi I2, An. maculatus, An. farauti, An. dirus, An. sinensis, An. atroparvus, An. darlingi, An. albimanus*. A description of the methods used to create the alignment is found in Neafsey *et al*. 2015. Phase 1 of the Ag1000G project comprises 590 *An. gambiae* and 129 *An. coluzzii* collected from 9 countries in Africa, plus 46 hybrid individuals from Guinea Bissau. A detailed description of the samples and data is given in Anopheles gambiae 1000 Genomes Consortium 2017.

### Identifying UCEs

To identify invariant regions we used only parts of the multi-species alignment where sequence data was available for all 21 strains. We used Variscan v2.03 (Vilella *et al*. 2005) to find regions of the alignment of 18bp or longer containing no variation. We mapped the resulting regions back to the PEST reference genome using BWA-aln with strict mapping parameters (exact parameters can be provided on request; bwa-0.7.10 (Li and Durbin 2010)). Sequences that mapped at multiple places in the genome were included in the analysis, but flagged as ‘repeat sequences’ as these would not be suitable for use as CRISPR targets. We used BEDTools (Quinlan and Hall 2010) to classify the genomic location of the UCEs (such as exonic, intronic etc). The AgamP4.12 basefeatures file was used from VectorBase (Giraldo-Calderón *et al*. 2015). Genic sequences were defined as those with an AGAP gene annotation so include exons, UTRs and introns. UCEs that partly or wholly fell within genes were classified by us as genic, and those outside genes were classified as intergenic.

For comparison, we used the same method to identify invariant sequences of 18bp or more just in the *An. gambiae* complex species (*An. gambiae PEST, An. gambiae s.s., An. coluzzii, An. merus, An. arabiensis, An. quadriannulatus, An. melas*). We also looked for conservation beyond the *Anopheles* genus by performing a BLAST with default search parameters (Altschul *et al*. 1990) search of the UCEs against *Culex quinquefasciatus* and *Aedes aegypti* reference genomes. From the BLAST results we extracted sequences of 18bp or more with no substitutions, insertions or deletions.

### Random control sequences

So that we could compare the location of UCEs with non-UCEs we extracted 10 randomly distributed sets of control sequences from the multi-species alignment file that were matched to give the same number of sequences with the same base-lengths. To compare variation in the Ag1000G data in UCEs and non-UCEs, we also extracted 10 sets of control sequences from the AgamP4 genome but also matching for genic and intergenic locations.

### Orthology between species

For UCEs that fell within genes, we compared the orthology identifiers between AgamP4 and *An. arabiensis* Dongola references genomes, and between *An. gambiae* PEST and *An. funestus* FUMOZ reference genomes. We chose these species because *An. gambiae* (and its sister species *An. coluzzii*), *An. arabiensis* and *An. funestus* are the most important malaria vectors in sub-Saharan Africa. *An. gambiae* PEST is a hybrid strain of *An. gambiae* and *An. coluzzii* (previously known as S and M forms of *An. gambiae*). *An. gambiae* and *An. arabiensis* are closely related (in the same species complex) and *An. funestus* is more distantly related. Genic UCEs were checked for orthology between *An. gambiae* and *An. arabiensis* and between *An. gambiae* and *An. funestus*. Coordinates of UCEs were extracted from the multiple-alignment file for *An. arabiensis* and *An. funestus* reference genomes, and annotated with gene names from the basefeatures files Anopheles-arabiensis-Dongola_BASEFEATURES_AaraD1and Anopheles-funestus-FUMOZ_BASEFEATURES_AfunF1.3 (from VectorBase). Orthology identifiers for each gene in each species were found from the ODBMOZ2_Anophelinae database at OrthoDb.org (Kriventseva *et al*. 2019). Orthology identifiers that match between species indicated that the genes were orthologous. We could not use orthology to directly compare intergenic UCEs, so instead we identified flanking genes for each intergenic UCE in the reference genome of each species, and then compared the orthology identifiers for these genes as before.

### Ontology analysis of genes containing UCEs

PANTHER software (version 14.0) (Mi *et al*. 2016) was used to categorise the gene ontology (GO-Slim) terms of the genes containing UCEs. A gene was represented in the analysis once, regardless of how many UCEs it contained. The genes were clustered by GO-Slim molecular function, biological process and cellular component terms.

Because the Panther categorisation does not take into account how much of the genome is covered by each GO term, we used GOseq (Young *et al*. 2010) to carry out length-bias corrected gene ontology (GO) enrichment analysis, implemented in Galaxy (Afgan *et al*. 2018). GOseq corrects for gene length using a Wallenius non-central hyper-geometric distribution. We used GO-Slim terms extracted from VectorBase (Giraldo-Calderón *et al*. 2015) for AgamP4.12 gene set. GO terms with a Benjamini-Hochberg corrected false discovery rate (FDR) of less than or equal to 0.05 were considered over-represented. We also looked for over-representation of GO-Slim terms in the genes flanking integenic UCEs. We were interested to see how our set of UCEs compared to UCEs from *Drosophila* studies, so as well as our full data set, we also performed the GO term analysis on a subset of genes that contained at least one UCE over 50bp long, to make the data comparable.

### Targets for mosquito control

One form of gene drive aimed at population suppression looks to disrupt essential mosquito genes and thereby impose a strong reproductive load on the population as it spreads. UCEs may offer good targets for control of *An. gambiae* by a gene drive method; if any sequence variation at these sites results in high fitness costs there would be little selective advantage to a mosquito having the variant allele over the gene drive allele. We searched the functional annotations of genes containing UCEs to find genes that may have a suitable function to be targeted for control. Gene descriptions were obtained from VectorBase (Giraldo-Calderón *et al*. 2015). Gene drives that confer recessive female sterility are particularly potent since both sexes can transmit the drive at very high rates to offspring yet only females homozygous for the drive display the phenotype, which results in a drastic reduction of the population’s reproductive capacity (Burt 2003, Burt and Deredec 2008). P-sterile values were available for some genes from (Hammond *et al*. 2016). P-sterile is a sterility index based on a logistic regression model that correlates gene expression features in *Anopheles* with the likelihood that mutations of the gene produce female sterile alleles in the model dipteran *Drosophila melanogaster* (Baker *et al*. 2011).

To narrow down the gene list to potential vector control targets, we leveraged the large amount of phenotype data already available for *Drosophila* mutants. Where possible, *Drosophila* orthologues were identified for genes containing UCEs (in Vectorbase). We used ID converter in FlyBase (Gramates *et al*. 2017) to batch convert *Drosophila* gene identifiers into alleles associated with the genes (FBal numbers). The alleles have associated phenotype data provided by the research community; we searched for phenotypes conferring female sterility or recessive lethality.

### Transcription factor binding site motifs in UCEs

We used the ‘Find Individual Motif Occurrences’ (FIMO, Grant *et al*. 2011) scanning module (MEME suite 4.12.0, Bailey *et al*. 2009) to look for transcription factor binding motifs in UCEs and controls. The UCEs were scanned for known insect transcription factor binding sites using weighted matrices from the JASPER CORE collection (Insect position frequency matrices 8th release (2020), Khan *et al*. 2018). The results were filtered by q-value to account for multiple tests. A cut-off of q<0.05 was used.

### Variation at UCE locations in Ag1000G data

Using the final filtered variant file from phase 2 of the Ag1000G project (described at https://www.malariagen.net/data/ag1000g-phase-2-ar1) we extracted single nucleotide polymorphisms for the UCEs identified above, and for matched non-UCE regions. Diversity statistics were calculated in scikit-allel v1.3.2 (Miles et al 2020): number of segregating sites (s), nucleotide diversity (pi) and the neutrality test Tajima’s D (Tajima 1989).

Data used in this study are publicly available from the *Anopheles* 16 genomes consortium and the *Anopheles* gambiae 1000 Genomes project. Data generated in this study are given in the supplementary tables.

## RESULTS

### Ultra-conserved regions from the multi-species alignment

Much of the MAF file does not include alignments of all 21 species and strains (see Table S8 in Neafsey *et al*. 2015). The total number of aligned bases from which we extracted the UCEs was 17,095,206 (7.4% of the AgamP4 reference genome (Suppl Table 1). A total of 8338 invariant regions of 18bp or more were identified; 1675 on chromosome arm 2L, 3015 on chromosome arm 2R, 1375 on chromosome arm 3L, 2188 on chromosome arm 3R and 85 on chromosome X (Table 1; we have also included the same metrics at different evolutionary timescales for comparison). The longest UCE was 164bp. Genomic coordinates of the UCEs relative to the *Anopheles gambiae* PEST reference genome are given in Supplementary table 2. The UCEs were distributed throughout the chromosomes, but were significantly under-represented on the X chromosome (See Supplementary figure 1 and Supplementary Table 1). The X chromosome is already under-represented in the MAF as it was less alignable than other chromosomes (see Figure 2 in Neafsey et al 2015). It is well established that the X chromosome shows higher differentiation between species than autosomes (due to ‘Haldanes Rule’ and the ‘Large X effect’) and genomic studies have reinforced this observation (Presgraves 2018). However, the under-representation in the MAF is not sufficient to explain the paucity of UCEs on the X. In the Anopheles genus, the X chromosome was observed to have undergone particularly dynamic evolution, with chromosome rearrangements at a rate of 2.7 times higher than the autosomes, and a significant degree of observed gene movement from X to other chromosomes relative to Drosophila (Neafsey et al 2015). This dynamic evolution of the chromosome may explain why it would be less likely to contain functional sequences that require conservation at the nucleotide level.

**Figure 1.**
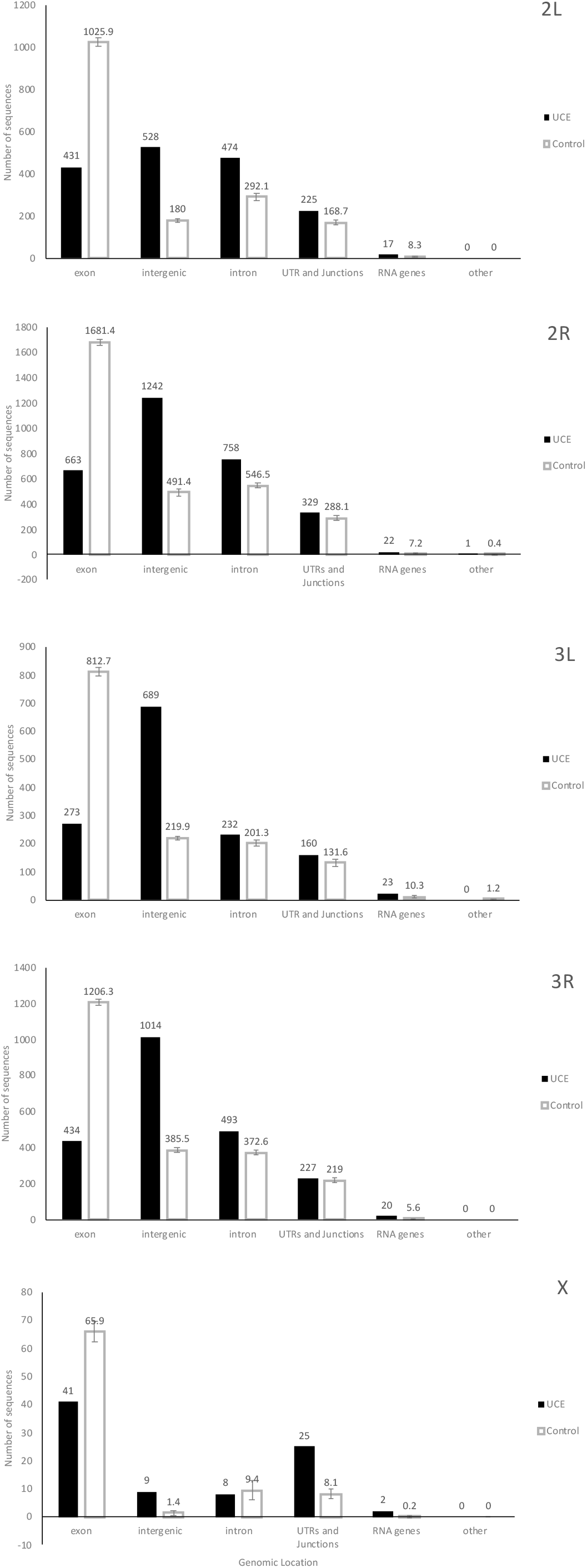
Distribution of UCE and non-UCE control sequences according to genomic location. Genomic locations annotated with BEDtools. Control error bars = standard deviation for 10 control data sets of sequences of matched length and number to the UCEs, extracted randomly from the MAF, only from regions where data for all 21 genomes is present.

**Figure 2.**
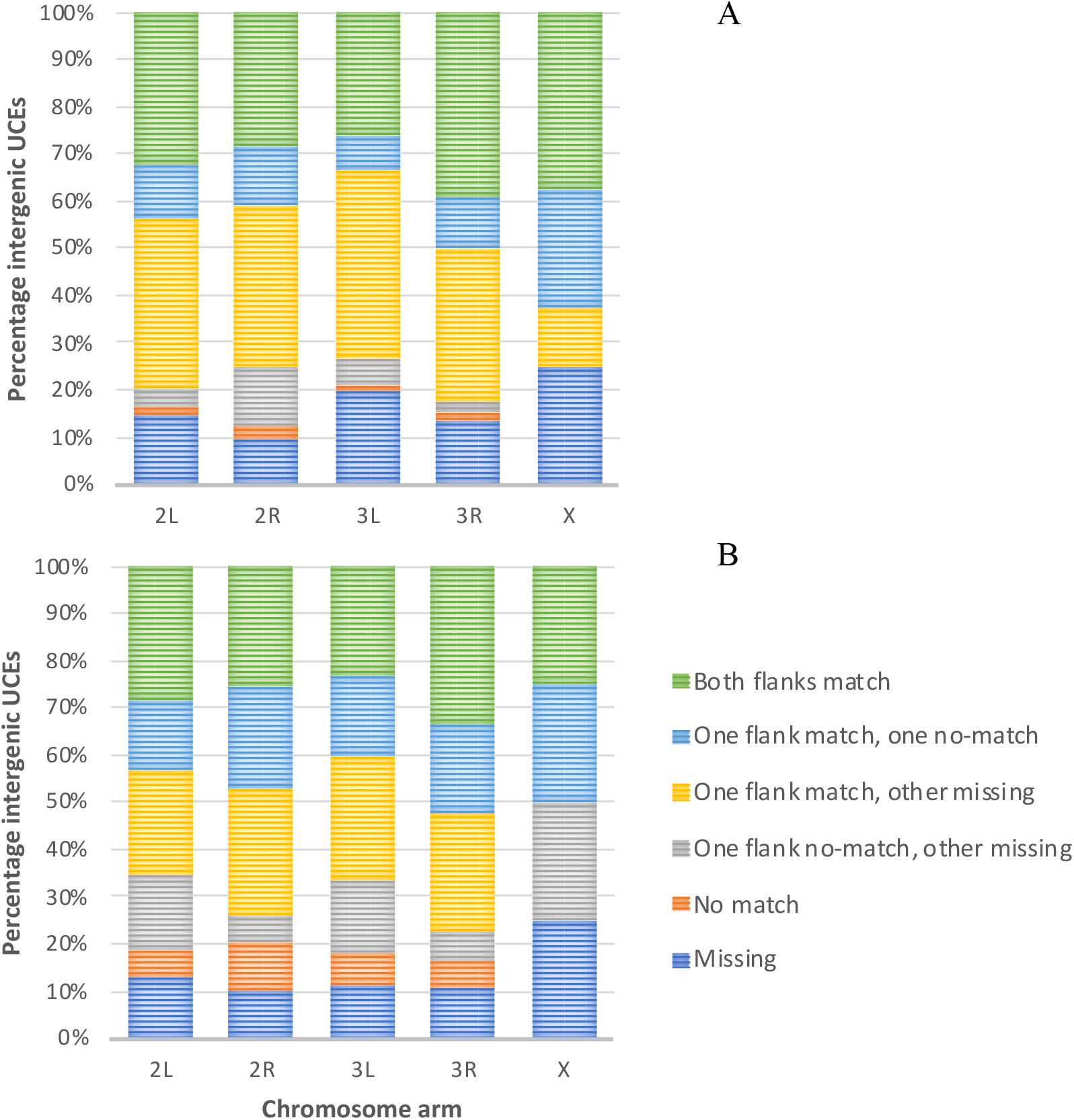
Number of intergenic UCEs that show synteny between A: *An. gambiae* and *An. arabiensis* and B: *An. gambiae* and *An. funestus*. The results are shown in six categories: matching orthology of both flanking genes, matching orthology of one flanking gene with no orthology on the other flank, matching orthology of one flanking gene with missing data on the other flank, no orthology on one flank with missing data on the other flank, no orthology of either flanking gene, and missing data on both flanks.

**Table 1.** Number of ultra-conserved sequences of 18bp or more, and total number of invariant sites within these sequences. *Gambiae* complex = 7 species and strains; *Anopheles* = 21 species and strains; *Culex* = *Culex quinquefasciatus* reference genome; *Aedes* = *Aedes aegypti* reference genome.

|  | 2L | 2R | 3L | 3R | X |
| --- | --- | --- | --- | --- | --- |
| *Gambiae* complex |  |  |  |  |  |
| No. UCEs | 452,281 | 612,824 | 376,383 | 498,473 | 99,561 |
| No. Invariant bases |  |  |  |  |  |
| within UCEs | 15,365,491 | 21,350,270 | 12,886,437 | 17,278,830 | 3,338,454 |
| *Anopheles* |  |  |  |  |  |
| No. UCEs | 1,675 | 3,015 | 1,375 | 2,188 | 85 |
| No. Invariant bases |  |  |  |  |  |
| within UCEs | 45,916 | 81,186 | 37,102 | 59,055 | 2,299 |
| *Anopheles+Aedes* |  |  |  |  |  |
| No. UCEs | 278 | 344 | 193 | 293 | 15 |
| No. Invariant bases |  |  |  |  |  |
| within UCEs | 8,161 | 10,275 | 5,499 | 8,339 | 456 |
| *Anopheles+Culex* |  |  |  |  |  |
| No. UCEs | 279 | 350 | 202 | 310 | 16 |
| No. invariant bases |  |  |  |  |  |
| within UCEs | 8,201 | 10,184 | 5,716 | 8,691 | 503 |
| *Anopheles+Aedes+Culex* |  |  |  |  |  |
| No. UCEs | 192 | 247 | 133 | 217 | 12 |
| No. invariant bases |  |  |  |  |  |
| within UCEs | 5,995 | 7,579 | 3,989 | 6,391 | 393 |

Size distributions of the UCEs are shown in Supplementary figure 2. In the autosomal genic UCEs there is pattern of a jump in frequency every 3 bases, indicating the tendency for runs of ultra-conserved bases to neither start nor end on third codon positions in coding regions. UCEs are significantly more AT-rich than random sequences (64% and 54% respectively, t-test p<0.001).

We annotated the UCEs in BEDtools to identify where they were found in the genome with regards to exons, introns, UTRs, intergenic regions etc (Figure 1). The 21-genome aligned parts of the MAF file from which we extracted the UCEs is not representative of the reference genome with respect to these features, so we extracted randomly distributed sets of ‘control’ sequences from the MAF, and only from sequences where all 21 genomes were aligned. These control sequences were matched to give the same number of sequences with the same base-lengths as the UCEs, and were compared with the UCE locations to see whether the UCEs were randomly distributed. The UCE sequences were significantly over-represented (compared to control sequences) in intergenic regions (42% vs. 15%, ANOVA, p<0.05) and in RNA genes (1% vs. 0.4%, p<0.05), and less frequent in exons (22% vs. 57%, p<0.05). The MAF itself is heavily skewed to including exonic sequences, as only about 7% of the *An. gambiae* genome as a whole is exonic (Holt et al. 2002).

### Orthology between important vector species

To ensure that the UCEs were not falling in sequences that had aligned across the 21 genomes by chance, we checked for orthology between some species in the UCEs. For UCEs that fell within genes this was done by simply by comparing orthology identifiers (from OrthDB.org) between *An. gambiae* and *An. arabiensis*, and between *An. gambiae* and *An. funestus*. For *An. gambiae* and *An. arabiensis*, 94% of autosomal genes containing UCEs shared orthology. For *An. gambiae* and *An. funestus*, this number was 87%. The amount of orthology fell to 54% and 63% for genes containing UCEs on the X chromosome. For UCEs that were intergenic, we looked at orthology of the flanking genes. The results fell into six categories: orthology of both flanking genes, orthology of one flanking gene with no orthology on the other flank, orthology of one flanking gene with missing data on the other flank, no orthology on one flank with missing data on the other flank, missing data on both flanks, and no orthology of either flanking gene. Ignoring missing data, 92% of intergenic UCEs showed full or half synteny between *An. gambiae* and *An. arabiensis*, and 77% of UCEs showed full or half synteny between *An. gambiae* and *An. funestus* (Figure 2).

### Functional profile analysis of the genes containing UCEs via GO term enrichment

Of the 13,796 genes annotated in the *Anopheles gambiae* PEST gene set Agam4.12, 1,601 (12.9%) had at least one UCE. We clustered the genes based on GO-Slim terms for molecular function, biological process and cellular component (Suppl Figure 3).

Because the clustering does not take into account the amount of the genome covered by each GO class, we carried out length-bias corrected GO term enrichment analysis. This showed that certain functional groups were over-represented compared with the whole *Anopheles* PEST reference gene set (Figure 3).

**Figure 3.**
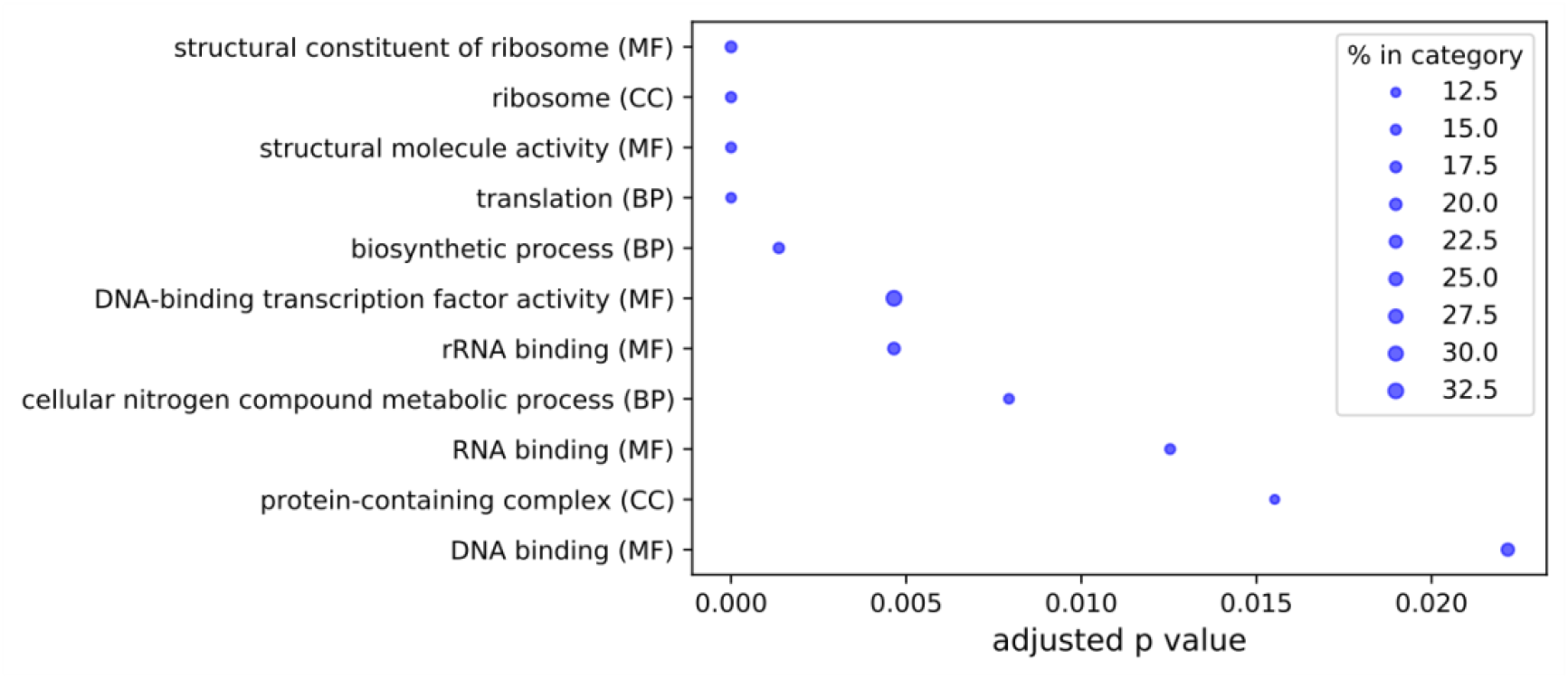
GOseq GO term enrichment analysis with length-bias correction. GO-Slim categories were extracted from the AgamP4.12 gene set. Results are shown for categories that were enriched with an FDR adjusted p-value<0.05. Bubble size is relative to the percentage of AgamP4.12 genes in a GO category that were present in the UCE gene set. MF=molecular function; BP=biological process; CC=cellular component.

In the genes containing UCEs over 50bp long, only 4 categories were over-represented: transmembrane transporter activity (MF), transmembrane transport (BP), transport (BP) and protein-containing complex (CC), (adjusted p values 0.0047, 0.0047, 0.0272, 0.0272 respectively).

Genes flanking intergenic UCEs were enriched for the GO-Slim categories DNA binding (MF), DNA-binding transcription factor activity (MF) and anatomical structure development (BP) (adjusted p values 4.16E-06, 1.46E-05 and 0.016 respectively).

### Potential targets for vector control

AGAP001189 (odorant-binding protein 10) contained the highest number of invariant bases in UCEs (1215/135306). Nine genes contained UCEs longer than 100bp, of which 3 are annotated as being involved in ion transport. These include the voltage gated sodium channel gene (VGSC, AGAP004707), which is a target for (and therefore has a significant role in conferring resistance to) some of the main classes of insecticides used for malaria vector control. VGSC is one of the most conserved genes we found, containing 13 UCEs with a total of 507 invariant bases, of which 91% were in exons and most coded for trans-membrane domains. A total of 357 genes contained 100 or more invariant bases. A full list of genes containing UCEs is given in Supplementary table 3.

Eleven genes containing UCEs had a p-sterile score of greater than 0.5 implying that they could be good targets to affect female fertility.

*Drosophila* orthologues were identified for 1309 of the 1601 genes containing UCEs. Allele and phenotype classes for these genes were extracted from Flybase where available. For an effective population suppression gene-drive, the target would affect female fertility or impose a genetic load as a homozygote, so we extracted UCE containing genes that have *Drosophila* orthologues annotated with a female sterile term or a lethal recessive term (shown in Supplementary table 3). 177 genes containing UCEs have *Drosophila* orthologues with an allele phenotype affecting female fertility, and 367 genes have *Drosophila* orthologues with an allele conferring a lethal recessive phenotype.

### Transcription factor binding motifs in UCEs

DNA binding motifs recognised by transcription factors might be expected to be constrained and hence enriched for UCEs since this protein:DNA interaction is sequence-specific. The FIMO search found that 38% of UCEs contained hits for insect transcription factor binding sites with a q value <0.05 (48% of intergenic and 30% of genic UCEs). This compared with 23% for control (non-conserved) sequences of the same number and length. On the X chromosome, where data is sparse (only 8 intergenic and 75 genic UCEs (Figure 4)), the numbers of transcription factors binding sites were not significantly different between UCEs and controls.

**Figure 4.**
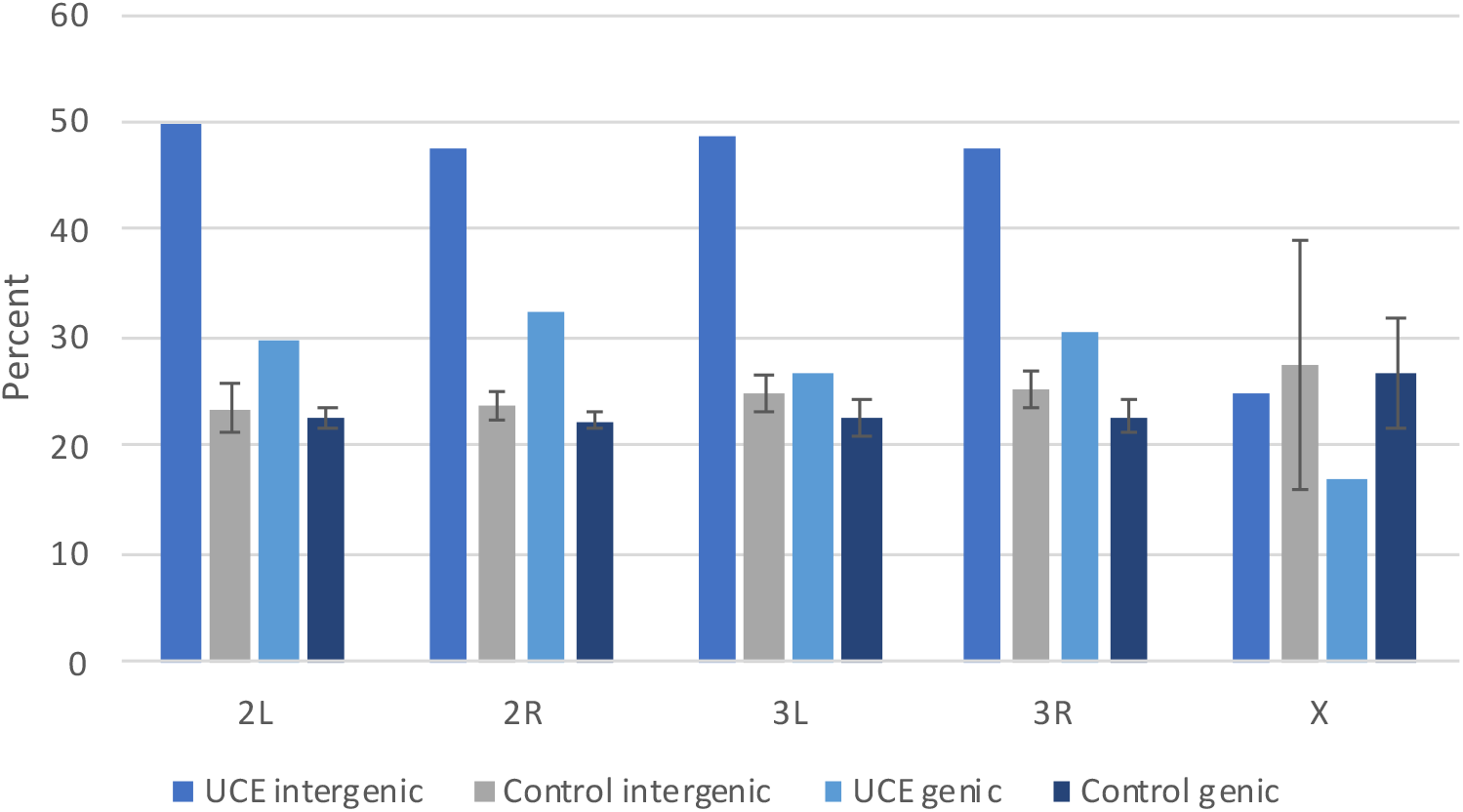
Percent of UCEs and control sequences that contain at least one insect transcription factor binding motif. Control error bars = standard deviation for 10 control data sets. UCEs were searched for known insect transcription factor binding sites from the JASPER CORE collection (Insect position frequency matrices 8th release (2020), Khan *et al*. 2018). The results were filtered by q-value to account for multiple tests. A cut-off of q<0.05 was used.

### Genetic variation at UCE locations in Ag1000G data

In order to see whether sequences that are ultra-conserved across the *Anopheles* genus show variation in wild mosquito populations, we searched for single nucleotide polymorphisms (SNPs) in the 1142 samples from phase 2 of the Ag1000G project. There were significantly fewer sites containing polymorphisms in UCEs than control sequences (Figure 5 middle), and those SNPs that were present were at significantly lower frequency (Figure 5 top). Of the 8338 UCEs, 1213 (15%) contained no SNPs in the 1142 samples (229 on 2L, 470 on 2R, 226 on 3L, 259 on 3R and 29 on X). Tajima’s D is significantly more negative for UCEs than controls, with the exception of X chromosome intergenic sequences (Figure 5 bottom). Negative values of Tajima’s D are expected for sequences under purifying selection.

**Figure 5.**
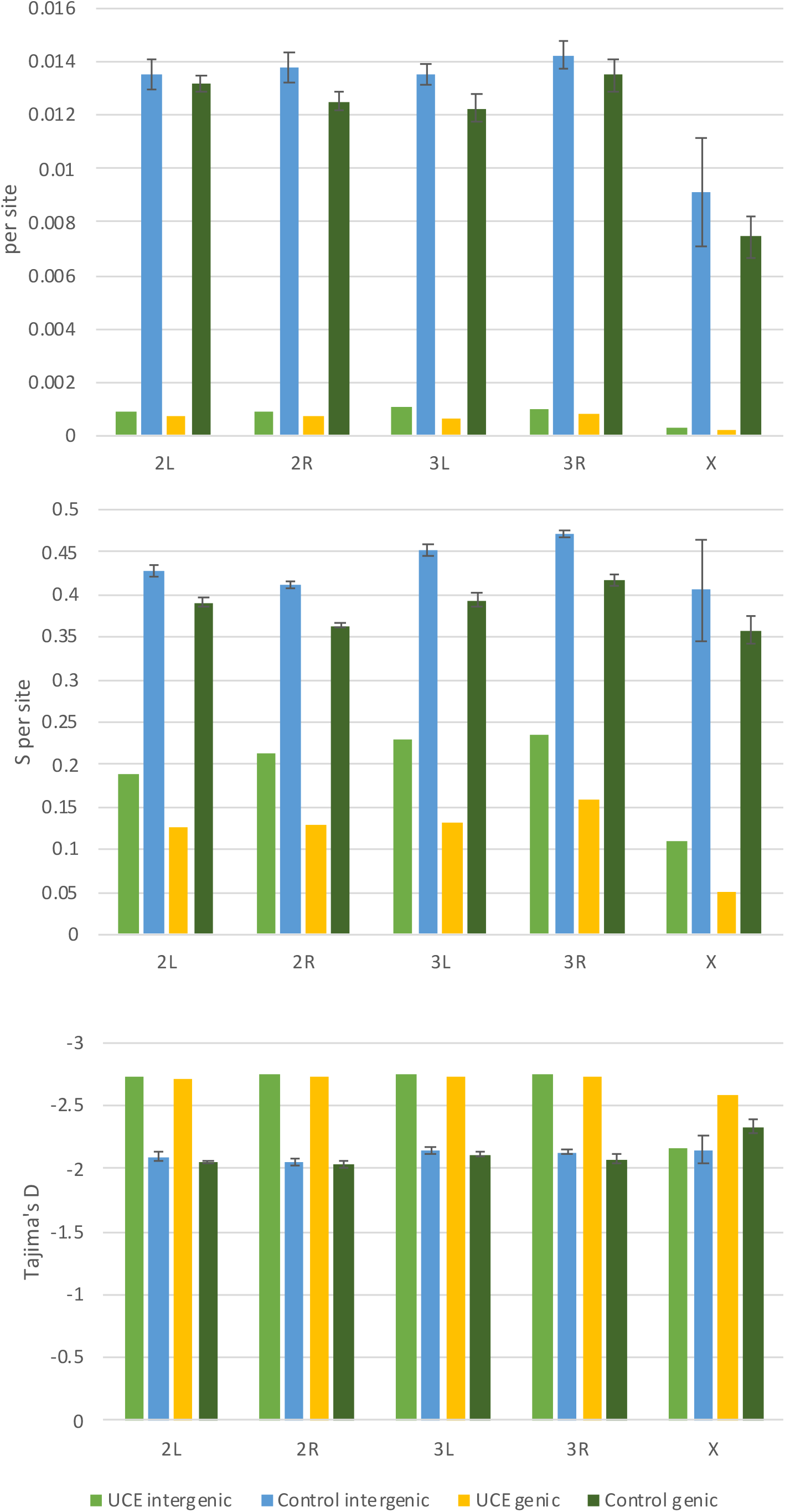
Genetic diversity per chromosome arm in 1142 *Anopheles gambiae s.l*. samples in UCE locations. Top: nucleotide diversity (π); middle: segregating sites (s); bottom: Tajima’s D. Calculations were made in scikit-allel v1.3.2 (Miles et al 2020).

The Ag1000G study (*Anopheles gambiae* 1000 genomes consortium 2017), performed a search within the Phase 1 data to look for potential Cas9 targets (non-overlapping exonic invariant sequences of 21bp, ending in the ‘NGG” motif). They identified 13 genes containing sequences corresponding to this motif. None of these 13 genes contained UCEs as defined by our study, so these genes also did not overlap with invariant sequences in Ag1000G data found here. We did not confine our search for UCEs to current Cas9 target site restrictions because of the growing possibility of relaxation of these constraints. However, for completeness we looked within our final set of UCEs for the Cas9 motif (18bp followed by −NGG, or CC-followed by 18bp). We found 1997 (24%) UCEs contained suitable targets for Cas9.

## DISCUSSION

### Similarities and differences of Anopheles UCEs with UCEs from *Drosophila*

Despite approximately 100 million years since their most recent common ancestor, we identified in the Anopheles genus over 8000 sequences of 18bp or more where there was no nucleotide variation across the alignment of 21 species and strains. By coincidence, this is approximately the same span of evolutionary time covered in the human/mouse/rat data set in which UCEs were originally identified (Bejerano *et al*. 2004). 481 UCEs of more than 200bp were observed between these genomes, but the longest we found in the *Anopheles* genus was 164bp. This is consistent with previous reports that UCEs are fewer and shorter in insects (mainly *Drosophila*) than vertebrates (Makunin *et al*. 2013; Glazov *et al*. 2005). Our criteria for identifying UCEs were somewhat different than those used previously. First, we only considered sequences that were present in all 21 species/strains in the alignment; some of these species have poorly assembled genomes, so this may have reduced the number of UCEs that we uncovered. Second, we also included invariant stretches of 18bp or more, whereas *Drosophila* studies have used cut-offs of 50bp (Glazov *et al*. 2005, Warnefors *et al*. 2016), 80bp (Kern *et al*. 2015) or 100bp (Makunin *et al*. 2013). Despite this we see some similarities between our UCEs and UCEs found in *Drosophila*. UCEs are located in all parts of the genome and, like *Drosophila*, the majority are found in intergenic regions and introns. We also found that junction locations (e.g. intron-exon, exon-intergenic etc) are over-represented compared to random sequences, which in *Drosophila* has been linked to conservation of splice-sites (Glazov *et al*. 2005; Warnefors *et al*. 2016). Another similarity with *Drosophila* is the high proportion of genes with the GO terms ‘binding’ and ‘transporter activity’ (Kern *et al*. 2015; Glazov *et al*. 2005). In *Drosophila*, ion channel/transporter genes have been shown to undergo extensive RNA editing (Hanrahan *et al*. 2000; Hoopengardner *et al*. 2003; Rodriguez *et al*. 2012) which is thought to explain the high level of conservation. This is because RNA adenosine deaminases require double stranded RNA as a substrate, which means that there is likely to be strong selection at the nucleotide level. The high number of UCEs in *Anopheles* ion channel/transporter genes suggests a similar mechanism is responsible for the high conservation in the *Anopheles* genus. However, these genes are extremely long and are not over-represented in the UCE data when a length-bias corrected analysis is carried out in GOseq. In the GOseq analysis, the most over-represented molecular functions are mostly involved in binding or structure. Transcription factor binding, enzyme binding and nucleic acid binding have also been shown to be associated with ultra-conservation in both invertebrates and mammals (Bejerano *et al*. 2004; Glazov *et al*. 2005). A noteworthy addition to highly represented GO terms in *Anopheles* that has not been reported in *Drosophila*, is the category of ‘catalytic activity’ genes, although again, these were not over-represented when gene length was taken into account. When the GO term clustering was carried out on genes containing UCEs of 50bp or more in length, we found that the category reduced from 28% to 18% suggesting that these shorter ultra-conserved regions most likely code for a small number of key residues around an active site.

The high number of UCEs that we observe in intergenic regions and introns suggests that we have found numerous unannotated locations in the *Anopheles* PEST reference genome with putative regulatory functions. At least 70% were syntenic between *An. gambiae*/*An. arabiensis* and *An. gambiae*/*An. funestus* so the location of these highly conserved sequences is likely to be important. A GOseq analysis of the genes flanking these intergenic sequences showed significant over-representation of genes with DNA binding GO terms. Sequences that are ultra-conserved at the nucleotide level across a long evolutionary time have been shown to be linked to regulatory functions such as cis-regulation of genes (e.g. enhancers, insulators, silencers) and RNA genes (e.g. miRNA, snRNA), likely because of the sequence-specific nature of protein:nucleotide or nucleotide:nucleotide interactions. 19 of the 77 miRNA genes that are annotated in the *Anopheles* PEST genome were included in our set of UCEs (other miRNAs may contain ultra-conserved regions that did not meet our criteria). We also found known insect transcription binding factors in 48% of the intergenic UCEs.

### Polymorphisms in UCEs in *Anopheles* populations

All of the UCEs discovered from the alignment of the reference genomes of 21 *Anopheles* species were also found to be highly conserved in the sample of 1142 wild caught mosquitoes sequenced in phase 1 of Ag1000G. Although the majority of UCEs contained one or more polymorphisms, they were almost all rare. 1213 UCEs showed no polymorphisms at all in this sample. This does not rule out the existence of polymorphisms in the wild populations, but does imply that there may be strong constraint at a nucleotide level that means alteration of the sequence either naturally or by the action of a gene-drive may have a strong fitness cost. This would need to be tested experimentally as different levels of underlying functional constraint may have different fitness costs. For instance, deletion of certain ultra-conserved sequences in mice gave no discernible fitness cost (Ahituv *et al*. 2007), but a similar experiment in *Drosophila* showed promise, with 4 out of 11 UCEs with inserted transposons having a lethal recessive phenotype (Makunin *et al*. 2013). For a resistance-proof gene drive, selecting target sites that show high levels of conservation is a good starting point, but the targets would need to be tested under selection pressure to ensure that functional mutants do not arise.

### UCEs and vector control

UCEs occur within many genes that could have potential for vector control. Nearly 200 genes have *Drosophila* orthologues with an allele phenotype affecting female fertility, and over three hundred genes have *Drosophila* orthologues with an allele conferring a lethal recessive phenotype. These phenotypes could both be used for a population suppression strategy i.e. to reduce the numbers of mosquitoes to a level where malaria could no longer be transmitted (Deredec *et al*. 2011). More investigation would be needed to see whether disrupting the genes at the ultra-conserved loci gives the same phenotype in *Anopheles*. There are also genes that confer recessive phenotypes in *Drosophila* such as ‘flightless’ or ‘behaviour defective’ that could also be used for population suppression, or for a population modification type of strategy, where instead of reducing the mosquito population it is replaced by a strain that cannot transmit malaria (Carballar-Lejarazú *et al*. 2018). Precise targeting of sequences using CRISPR/Cas9 gene editing had made testing for these phenotypes feasible.

Another potential mode of mosquito genetic alteration that has not yet been explored would be to target sequences involved in gene regulation. Many ultra-conserved sequences in mammals and invertebrates are thought to be involved in regulation of genes important in development.

Targeting a sequence that is conserved between species means that the gene drive could spread between closely related species that hybridise in the wild. For this to happen the species would need to mate in the wild, produce some fertile offspring, and be able to express the CRISPR enzyme using the same promoter. Three species (*An. gambiae*, *An. coluzzii* and *An. arabiensis*) are responsible for the majority of malaria transmission in some parts of sub-Saharan Africa, and are known to hybridise in nature (e.g. Weetman *et al*. 2014, Anopheles gambiae 1000 Genomes Consortium 2017, Fontaine et al 2015). For effective vector control it would be desirable to be able to reduce or alter all three species with one construct. The gene drive would not spread to *Anopheles* species that do not mate in the wild, so would not spread beyond the *Anopheles gambiae* species complex. However, if a particular target site was proved to be effective for vector control in *An. gambiae*, a gene drive targeting an orthologous site could be developed in the laboratory for other important malaria vectors such as *An. funestus*.

## CONCLUSION

Thousands of short genomic regions exist that are conserved across the *Anopheles* genus. These sequences show many of the same traits as ultra-conserved elements found in *Drosophila* (such as an association with gene regulation and ion channel activity). Our list of ultra-conserved elements in the *Anopheles* genus should provide a valuable starting point for the selection and testing of new targets for gene-drive modification in the mosquitoes that transmit malaria. Focussing on sequences that have been tightly conserved over long evolutionary time has promise for mitigating against or slowing the development of resistant alleles in the wild population.

## Authors’ contributions

SO and AB jointly devised the study; SO performed and guided data analysis and wrote the manuscript; AF carried out data analysis; SF designed and performed preliminary GO term analysis; TD assisted in bioinformatics; TN and AC gave advice on analysis; all authors provided editorial comments and read and approved the final manuscript.

## Acknowledgements

Many thanks to the Anopheles 16 genomes consortium and MalariaGEN for permission to use the 21 species alignment data and the Ag1000G variation data. Thanks also to Howard Lewis for providing several custom data handling scripts.

