## Supplemental figures and table 1 for "Ultra-conserved sequences in the genomes of highly diverse *Anopheles* mosquitoes, with implications for malaria vector control"

| Chromosome arm | %AgamP4 aligned with 21 genomes in MAF* | %MAF** covered by UCEs | %AgamP4 covered by UCEs |
|---|---|---|---|
| 2L | 7.594 | 1.225 | 0.093 |
| 2R | 9.340 | 1.412 | 0.132 |
| 3L | 6.737 | 1.312 | 0.088 |
| 3R | 7.181 | 1.546 | 0.111 |
| X | 3.898 | 0.242 | 0.009 |
| Whole genome | 7.418 | 1.319 | 0.098 |

**Supplementary Table 1. MAF statistics.** *The MAF is formatted as blocks of aligned sequences, with most blocks containing fewer than 21 genomes. This is the proportion of AgamP4 in the MAF that was aligned with all 21 genomes. **MAF containing aligned sequences of all 21 reference genomes.

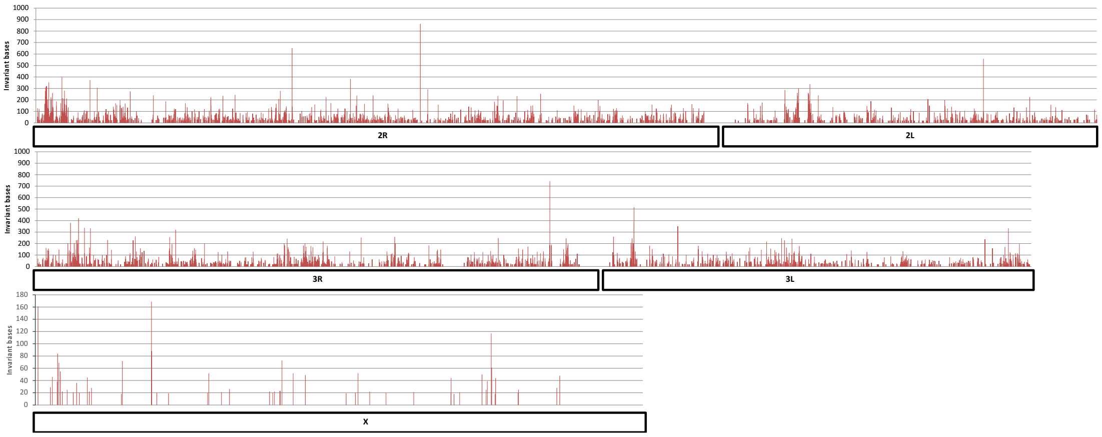


**Supplementary Figure 1. Total length of invariant bases within UCEs in non-overlapping 10kb windows across the *Anopheles gambiae* genome.**

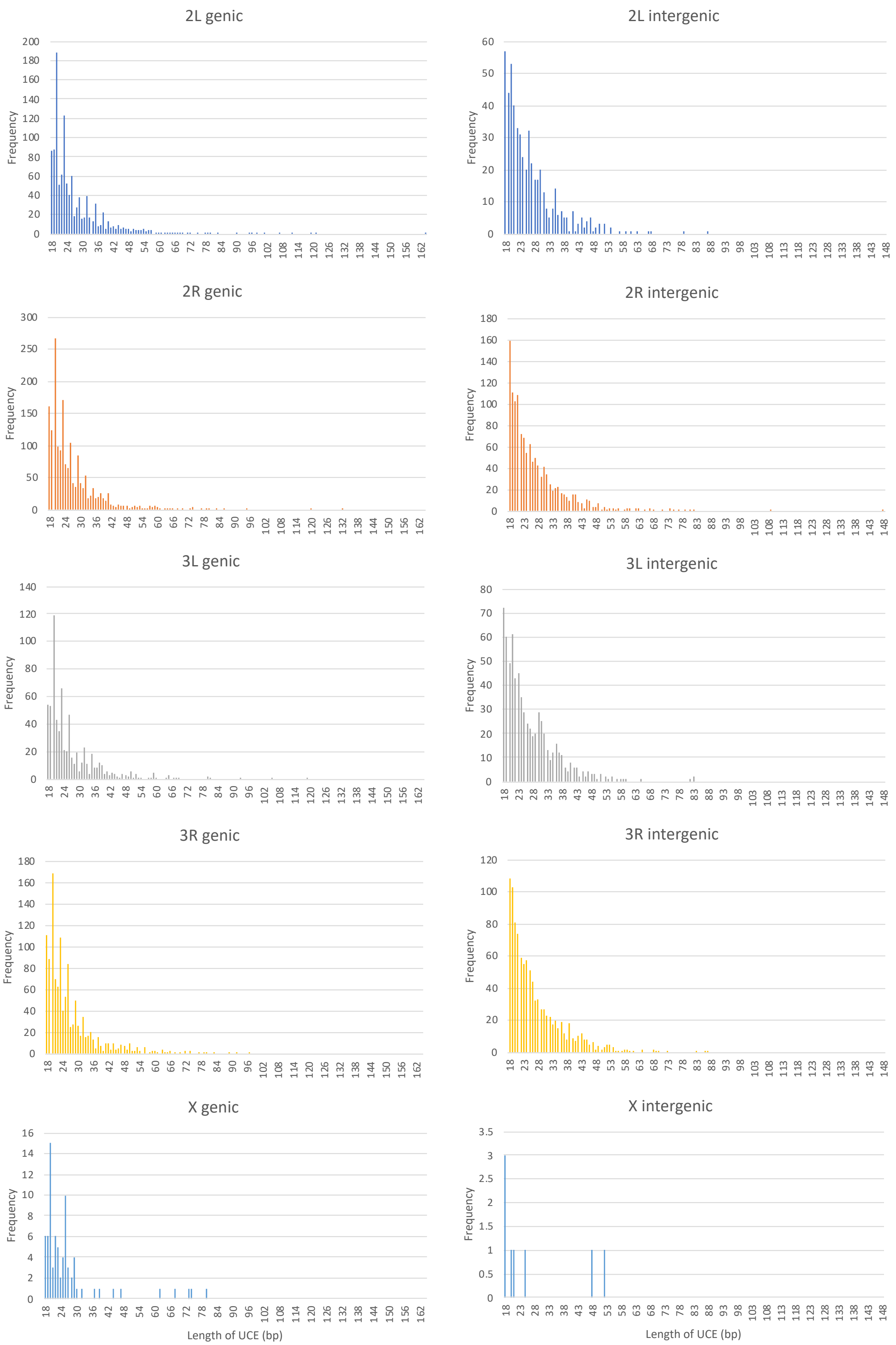
2L genic
2L intergenic
2R genic
2R intergenic
3L genic
3L intergenic
3R genic
3R intergenic
X genic
X intergenic
Frequency
Length of UCE (bp)

**Supplementary Figure 2. Distribution of sizes of UCEs.** Left panel: genic sequences; right panel intergenic sequences. Sequences containing any bases within a gene were treated as genic, all other sequences as intergenic.

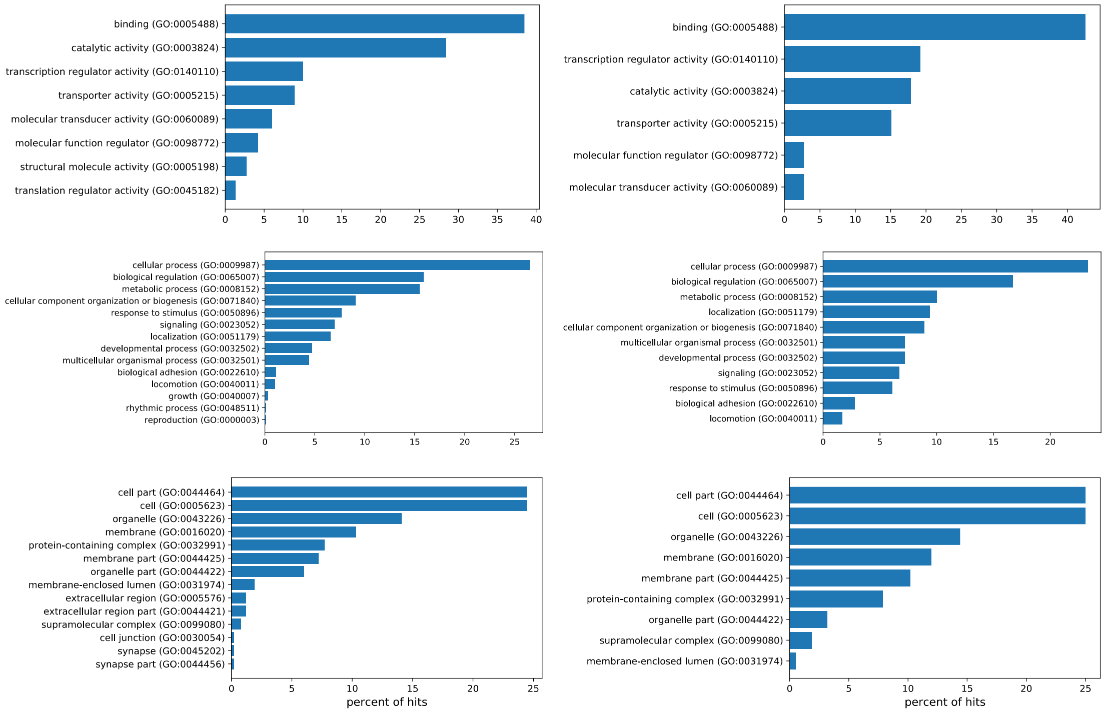


**Supplementary Figure 3. Panther clustering of GO terms by GO-Slim terms.** Top: molecular function; middles: biological process; bottom: cellular component. Left panel: genes containing UCEs; right panel: genes containing UCEs of length 50bp or over (for comparison with previous studies).
